# Mechanical Coordination of Intestinal Cell Extrusion by Supracellular 3D Force Patterns

**DOI:** 10.1101/2025.07.03.661686

**Authors:** Marija Matejčić, Meng Wang, Elia López Serrano, Carlos Pérez-González, Ronja Houtekamer, Gerardo Ceada, Pere Roca-Cusachs, Martijn Gloerich, Xavier Trepat

**Affiliations:** Institute for Bioengineering of Catalonia (IBEC), The Barcelona Institute of Science and Technology (BIST), Barcelona, Spain; Institut Curie, PSL Research University, CNRS UMR 144, Paris, France; Center for Molecular Medicine, University Medical Center Utrecht and Utrecht University, Utrecht, The Netherlands; IRTA, Programa de Sanitat Animal, Centre de Recerca en Sanitat Animal (CReSA), Campus de la Universitat Autònoma de Barcelona (UAB), Bellaterra, Barcelona, Spain; Unitat mixta d’Investigació IRTA-UAB en Sanitat Animal, Centre de Recerca en Sanitat Animal (CReSA), Campus de la Universitat Autònoma de Barcelona (UAB), Bellaterra, Barcelona, Spain; Department of Cell Biology, Physiology and Immunology, Universitat Autònoma de Barcelona, Bellaterra, Barcelona, Spain; Facultat de Medicina, Universitat de Barcelona, Barcelona, Spain; Institució Catalana de Recerca i Estudis Avançats (ICREA), Barcelona, Spain; Centro de Investigación Biomédica en Red en Bioingeniería, Biomateriales y Nanomedicina (CIBER-BBN), Barcelona, Spain

## Abstract

Every day, the mammalian intestinal epithelium extrudes millions of cells to sustain tissue self-renewal. Despite its fundamental role in intestinal homeostasis, the mechanisms that trigger, compartmentalize, and execute intestinal cell extrusion remain largely unknown. Here, using intestinal organoids, we map the three-dimensional forces and cytoskeletal dynamics that drive intestinal cell extrusion. We show that, unlike in other epithelia, extrusion is initiated by the sudden dissolution of a contractile myosin 2A meshwork triggered by a calcium influx. Following meshwork dissolution, the extruding cell and its neighbors generate an upwards traction force that requires myosin contractility but is generated by lamellipodial protrusions in neighboring cells. Importantly, these lamellipodia not only act as force generators but also determine whether extrusion occurs apically or basally, serving as symmetry breakers of the process. Finally, we show that compartmentalization of cell extrusion to the outside of the intestinal crypt does not require curvature and instead depends on myosin 2A. Our findings reveal that the intestinal epithelium exhibits a distinctive mode of extrusion, in which tension differentials—rather than compressive stresses from crowding—trigger and compartmentalize cell removal.

## Main Text

The mammalian intestinal epithelium is a rapidly self-renewing tissue that acts as a multifunctional barrier between the body and the external environment. A central mechanism enabling this renewal is cell extrusion—the removal of single cells from the epithelium without compromising barrier function. In the intestine, cell extrusion takes place in the differentiated villus compartment of the tissue. If dysregulated, excessive extrusion destroys barrier integrity and drives villus atrophy in precancerous pathologies like the inflammatory bowel disease^1–3^. Although the impact of extrusion on intestinal health is clear, nearly 80 years after the first description of the extensive ‘cell shedding’ in the intestinal villus^4^, much remains to be explored about how this process occurs within this complex epithelial environment.

Cell extrusion is a mechanical process (rev. in ^5,6^) that has been extensively investigated within simple epithelia such as MDCK cell lines. These studies, which typically injure a cell or otherwise promote cell death, found that key mechanical events of extrusion are spatiotemporally coordinated between the extruding cell and its neighbors^7–12^. A common feature is the formation of contractile actomyosin rings that act like a purse string to squeeze the extruding cell out of the tissue. Such rings have been observed both in the extruding cell^8,9^ and in its immediate neighbors^9,10,13^. In addition to actomyosin contractility, neighboring cells can contribute to extrusion by forming lamellipodial protrusions^10,14–16^, depending on cell viability^17^, the nature of the damage^16^ and tissue density^15^. The state of mechanical stress in the monolayer is a key determinant for homeostatic cell extrusion, with a consensus view that extrusion is promoted by compressive stresses and deferred by tensile ones^17–20^. Despite this emerging consensus, even in simplified systems such as immortalized cell line monolayers, we still lack measurements of key mechanical variables that drive extrusion—particularly the 3D physical forces responsible for expelling cells from the tissue.

The inaccessibility of the *in vivo* intestinal tissue has hindered mechanistic studies on intestinal cell extrusion. Work using fixed human samples showed that intestinal cells accumulate non-muscle myosin 2A during extrusion^21,22^. However, these snapshots lack temporal information of what is a highly dynamic process. This limitation is circumvented by intestinal organoid models, where actin was shown in live images to accumulate at the interface between the extruding cell and its immediate neighbors throughout extrusion^23^. Despite these advances, however, fundamental questions remain unanswered including the nature of the signals that trigger intestinal cell extrusion, the factors that restrict it to the villus region, and the mechanisms responsible for its execution. Here we address these questions by studying the 3D mechanics of cell extrusion in intestinal organoids. We show that extrusion is mainly compartmentalized to the villus by the activity of myosin 2A. Unlike previously studied systems, we demonstrate that intestinal cell extrusion is triggered by the collapse of a tensed actomyosin basal meshwork, triggered by a calcium influx. In addition, by mapping the 3D force patterns that drive cell extrusion, we show that neighboring lamellipodia are the main traction force generators in the system and act as symmetry breakers, determining whether cells extrude apically or basally. Together, our findings reveal a new mode of cell extrusion that, rather than being triggered by compression as traditionally recognized^17–20^, occurs under tension.

## Results

### Tissue curvature is not required to localize cell extrusion to the villus compartment of the intestine

To gain experimental access to the intestinal epithelium, we cultured organoids as monolayers on top of soft synthetic substrates^24^ (Fig. 1a). This tissue configuration preserves the crypt-villus compartments of the *in vivo* intestine but additionally enables live imaging with high spatial and temporal resolution. Unlike conventional 3D organoids, these open-lumen organoids present the apical lumen of the organoid as an open and accessible tissue surface (Fig. 1b), allowing us to image apical, lateral and basal cell surfaces (Video 1).

**Figure 1.**
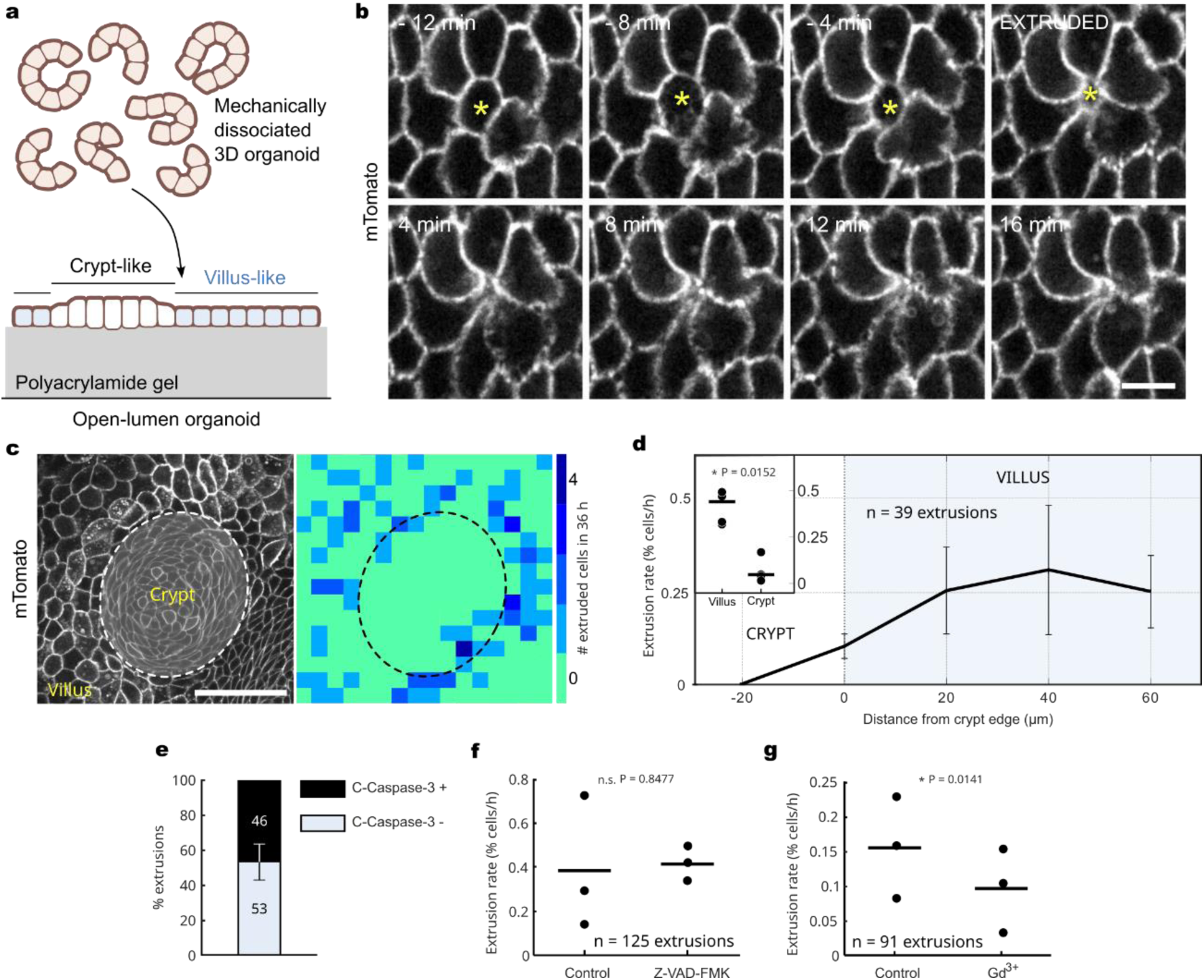
Tissue curvature is not required to localize intestinal cell extrusion to the villus. **a.** Schematic of the generation of open-lumen intestinal organoids grown atop a polyacrylamide hydrogel (grey). **b.** Spinning-disk timelapse of the extrusion of a villus cell (yellow asterisk). Cell membranes are labelled with membrane-tagged tdTomato (mTomato). Scale bar is 10 µm. **c.** (Left) Open-lumen mTomato monolayer image used to generate heatmap on the right. (Right) Heatmap of extrusion events marked over 36 hours of live imaging of the monolayer on the left at 4-minute time resolution. The outer edge of the intestinal crypt compartment is marked by the dashed line. Scale bar is 50 µm. **d.** Cell extrusion rate as a function of distance from the crypt edge. Data from 3 independent experiments. Inset: cumulative extrusion rate for the 3 experiments used for the main plot. **e.** Bar plot of fractions of positive and negative extruding cells in samples immunostained for cleaved caspase-3 (n = 3 experiments). Mean +/- s.d. Total number of counted extrusions is shown on each bar stack. **f.** Extrusion rate (% of cells extruding out of all cells, per hour) in open-lumen organoids treated with DMSO (0.1 %) or the pan-caspase inhibitor zVAD-FMK (50 µM). Data is from 3 independent experiments, 1 monolayer per experiment. **g.** Extrusion rate in DMSO or gadolinium (III) chloride (20 µM) treated open-lumen organoids, to block mechanosensitive ion channel function. Data is from 3 independent experiments, 1 monolayer per experiment.

In the *in vivo* intestinal epithelium, extrusion is restricted to the differentiated cells of the villus, predominantly at its tip^4,21,25^. To examine if our flat intestinal tissue captures this physiological feature, we mapped the extrusion distribution throughout the monolayer. As observed *in vivo*, cells in organoids extruded almost exclusively in the villus-like region containing differentiated cells (hereafter referred to as “villus” for simplicity, Fig. 1c). This indicates that the hallmark curvature *in vivo* is not required to localize cell extrusion to the intestinal villus. Moreover, we found that cells extruded uniformly throughout the villus region (Fig. 1d), additionally implying that neither cell age nor distance from the crypt triggers extrusion.

We next tested the long-held assumption that cells in the small intestine die before they extrude from the villus^4,21^. We examined the viability of extruding intestinal cells by staining for the apoptosis marker cleaved caspase-3. Only around 50% of the cells extruded from the organoid villus-like region were positive for cleaved caspase-3 (Fig. 1e), indicating that intestinal cells can extrude alive. Because of reports that different tissue preparation protocols might affect results on intestinal cell viability^21^, we also imaged living organoids with the fluorescent caspase 3/7 substrate NucView. Cells became positive for NucView fluorescence only after being fully extruded^23^, in some cases several hours after extrusion (Extended Data Fig. 1a), indicating that we were witnessing predominately live cell extrusion. Consistently, inhibiting apoptosis by a pan-caspase inhibitor zVAD-FMK did not reduce intestinal extrusion rate (Fig. 1f). By contrast, a reduction in extrusion rate was achieved by blocking mechanosensitive ion channels (Fig. 1g), known to be key upstream triggers of homeostatic cell extrusion^17^. Despite resulting in variable percentages of viability, these different approaches indicate that most small intestine cells in organoid monolayers extrude from the tissue while they are still living. Together, this initial investigation shows that intestinal cells do not require tissue curvature, stromal tissue, or apoptosis to extrude specifically in the intestinal villus.

### Upward and outward cell-substrate tractions characterize intestinal cell extrusion

The extrusion of cells from a tissue is a mechanical process that involves force generation in both the extruding cell and its neighbors^7–12^. To date, force measurements have been restricted to in-plane traction components, despite the fact that extrusion is fundamentally an out-of-plane process^20,26^. To fill this gap, we measured the time evolution of 3D cell-substrate traction fields generated during villus cell extrusion (Fig. 2ab, Video 2). The dominant force component was an upwards-pointing traction that peaked mid extrusion and decayed slowly over time, remaining significantly positive well after (>1 h) extrusion was completed. We also observed weaker but systematic in-plane traction components both under the extruding cell and its neighbors. To analyze the full temporal evolution of these traction fields, we represented both vertical and radial traction components as kymographs.

Vertical traction kymographs showed that the upwards-pulling traction generated under the extruding cell was balanced by downwards-pushing traction under its nearest neighbors (Fig. 2d, N1), resulting in the generation of a mechanical torque. Radial traction kymographs showed that, under the extruding cell, in-plane tractions pointed away from the extrusion center (Fig. 2e). Further away, across at least two rows of neighboring cells, in-plane tractions showed a broad signature in the opposite direction (Fig. 2e), suggesting a cooperative neighbor contraction towards the extrusion center.

**Figure 2.**
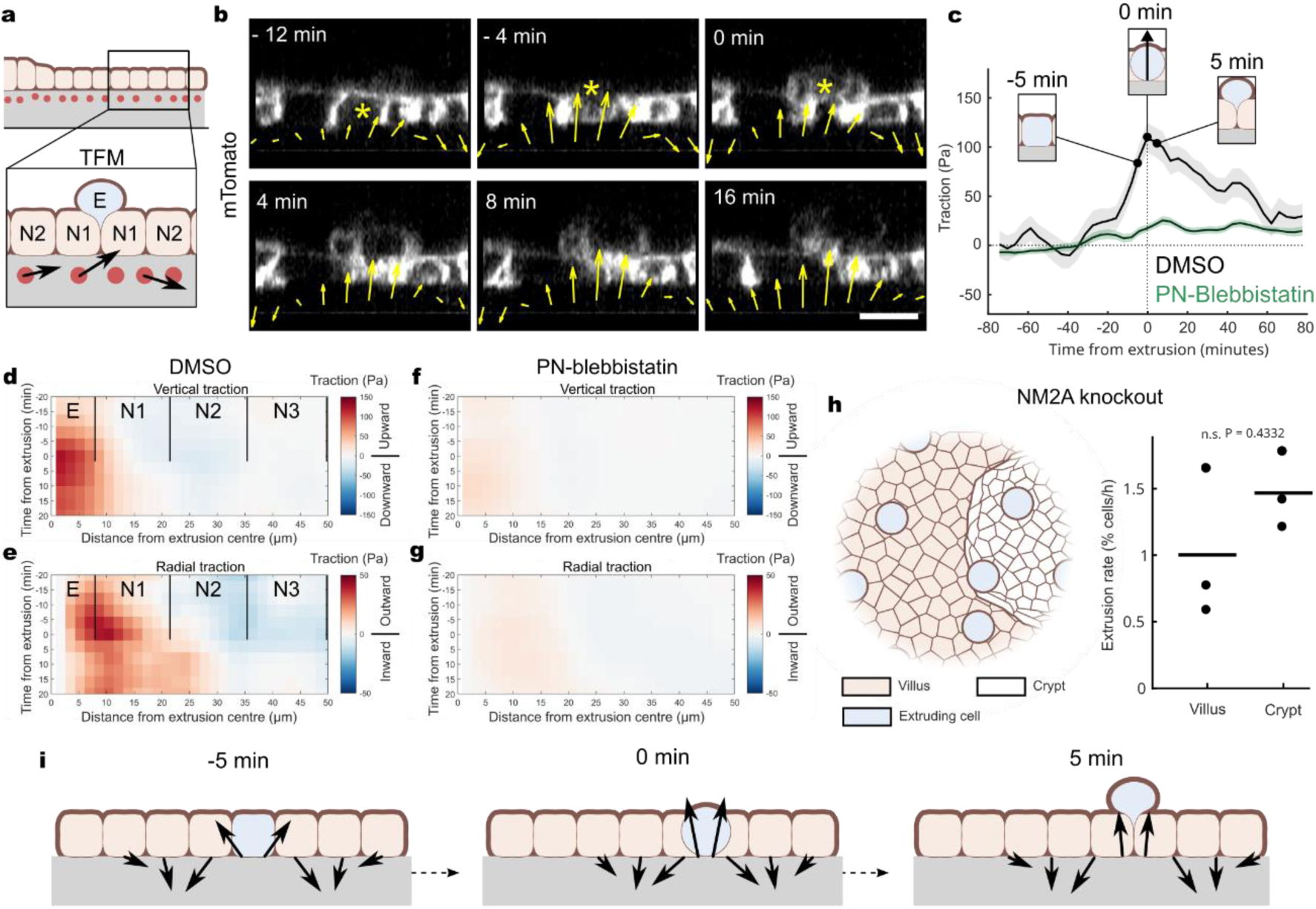
Supracellular 3D force patterns arise during cell extrusion. **a.** Schematic of a 2.5D traction force microscopy (TFM) experimental setup. Open-lumen organoids are grown on top of polyacrylamide hydrogels (grey) with fluorescent beads embedded near their surface. Displacement of beads signal is tracked in 3D and compared to a reference image to calculate the stresses that cells exert onto the substrate over time. E denotes the extruding cell, and Ni denotes the i^th^ neighbor cell. **b.** XZ-view of a spinning disk timelapse of an intestinal extrusion event (yellow asterisk) and traction vectors measured with TFM. Scale bar is 10 µm. **c.** Average out-of-plane traction under the extruding cell (6 µm radius around cell center) as a function of time from mid extrusion (mean +/- s.e.m.). Data from 44 extrusions (DMSO) and 59 extrusions (25 µM PN-blebbistatin), 3 independent experiments. Insets: illustration of the morphologically relevant timepoints of the extrusion process in 12 µm diameter-of-interest. **d.** Average kymograph of out-of-plane tractions in control open-lumen organoids, relative to the center of the extruding cell. Tractions are plotted as a function of distance from the center of the extruding cell, and of time from middle of the extrusion process. Data from 44 extrusions, 3 independent experiments. Same cells as in c (DMSO) and e. E denotes average area under the extruding cell, and Ni denotes area under i^th^ neighbor layer x. **e.** Average kymograph of in-plane tractions in the control open-lumen organoids as a function of distance from the center of the extruding cell, and of time from middle of the extrusion process. Same cells as in c (DMSO) and d. E denotes average area under the extruding cell, and Ni denotes area under neighbor layer i. Note the narrower range in the traction colormap scale compared in panel d. **f.** Average kymograph of out-of-plane tractions in the open-lumen organoids treated with 25 µm paranitro-blebbistatin. Plotted in the same way as d. Data from 59 extrusions, 3 independent experiments. **g.** Average kymograph of out-of-plane tractions in the open-lumen organoids treated with 25 µm paranitro-blebbistatin. Plotted in the same way as e. Same cells as in c and f. Note the narrower traction colormap range compared to panel f. **h.** Extrusion rate in villus and crypt compartments in NM2A knockout open-lumen organoids. Compare to wildtype data in inset in Fig. 1d. Data is from 3 independent experiments, 1 dish per experiment. **i.** Schematic summarizing 3D cell-substrate tractions under the extruding cell and its neighbors during intestinal cell extrusion.

Treatment with 25 µM paranitro-blebbistatin, used to inhibit non-muscle myosin 2A-generated cell contractility, did not stop extrusion events, but it reduced drastically both out-of-plane (Fig. 2f) and in-plane tractions (Fig. 2g). This result indicates that non-muscle myosin 2A (from now on ‘myosin 2A’) generates 3D forces during villus cell extrusion and that, when tension is removed from the monolayer, the forces involved during extrusion are markedly lower. Strikingly, with loss of contractility achieved either through paranitro-blebbistatin treatment or NM2A knockout, crypt cells begun to extrude at the same rate as villus cells (Fig. 2h). This latter finding indicates that myosin 2A-driven contractility prevents intestinal crypt cells from extruding.

Overall, our traction measurements reveal a supracellular 3D mechanical pattern during cell extrusion, whereby the extruding cell and its neighbors generate a mechanical torque, akin to the torque produced when sand is shoveled from a pile (Fig. 2i). As extrusion was not abolished with contractility inhibition, our results also indicate that the intestinal extrusion force adapts to the mechanical state of the villus tissue. Furthermore, we show that contractility of the crypt compartmentalizes intestinal cell extrusion exclusively to the villus.

### Intestinal cells extrude following the dissolution of a tensed basal myosin meshwork

Previous studies using cell lines have shown that extrusion is triggered by compressive stress^17–20^. However, our prior work showed that the villus domain in open-lumen organoids is under tension^24^, raising the question of how extrusion is triggered in this system. To address this question, we first characterized the distribution of myosin 2A during extrusion by imaging live organoids with endogenously tagged (NM2A-eGFP). In these organoids, we observed striking differences in the organization of basal myosin 2A between the villus and the crypt (Fig. 3a). In the crypt, myosin 2A was localized throughout the cell cortex (Fig. 3b, Extended Data Fig. 1b). By contrast, in villus cells, which contained overall higher levels of myosin 2A (Extended Data Fig. 1f, and ^27^), myosin 2A localized predominantly to a medial meshwork at the basal surface (Fig. 3bc), often forming a thick ring-like structure (Fig. 3d). Super-resolution Airyscan images revealed that, in this basal villus cell meshwork, myosin 2A formed regularly spaced clusters along basal stress fibers (Extended Data Fig. 1e). Unlike F-actin, myosin 2A was absent from villus cell–cell junctions (Fig. 3d, Extended Data Fig. 1c,d). A similar non-junctional myosin 2A structure has recently been imaged *in vivo*^27^, confirming that our open-lumen organoids recapitulate physiologically relevant mechanical features of the intestinal epithelium.

**Figure 3.**
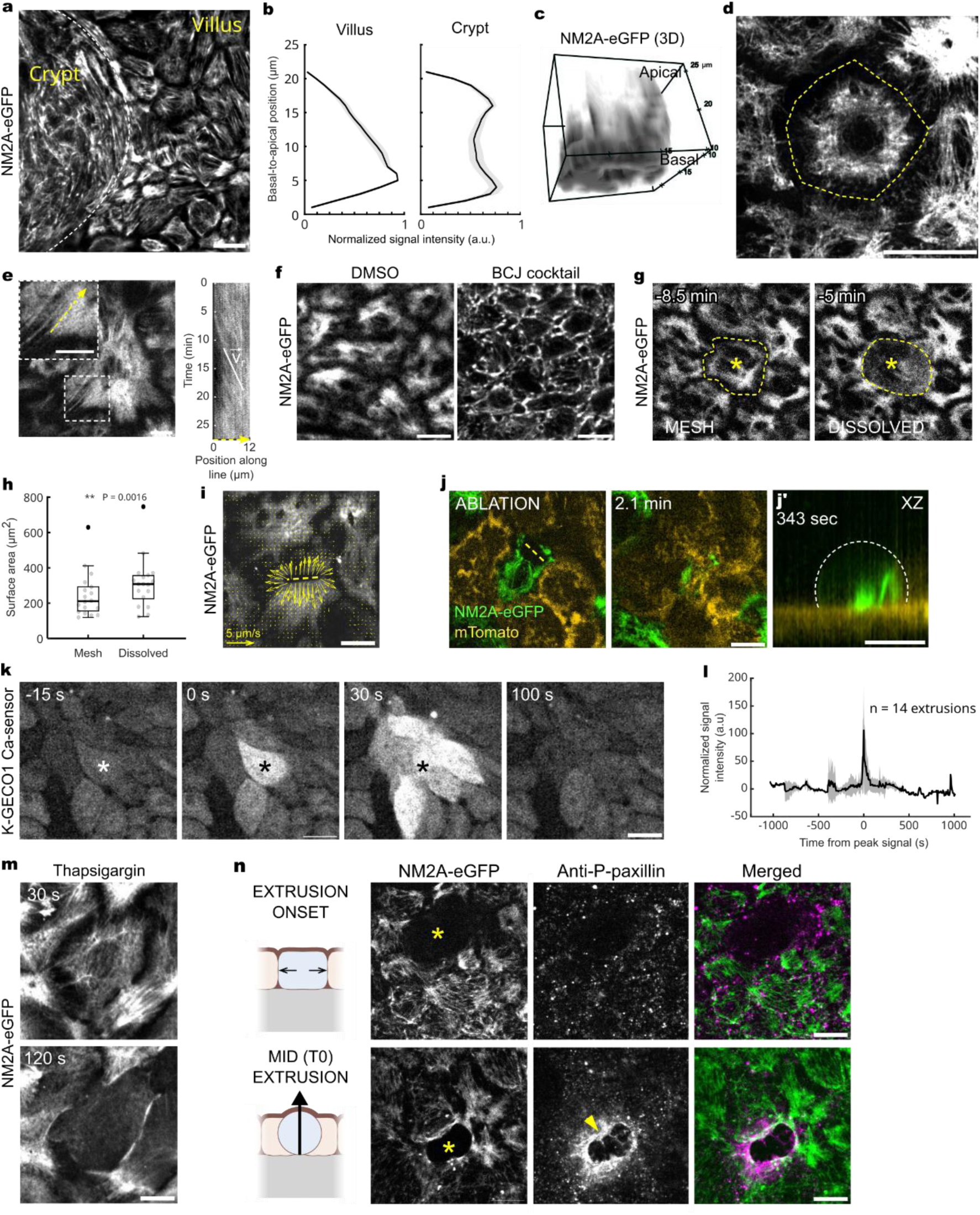
The dissolution of a tensed basal myosin meshwork initiates villus cell extrusion. **a.** Extrusion dynamics were imaged at the basal-most plane of the intestinal epithelium grown atop a polyacrylamide hydrogel. Spinning disk image of the fixed open-lumen organoid with the non-muscle myosin 2A (NM2A) heavy-chain protein tagged with GFP. Scale bar is 10 µm. **b.** Apico-basal distribution of NM2A-eGFP fluorescent signal in villus and crypt compartments, height is normalized for each cell to the average of the population. Data from 5 independent experiments, 1 monolayer per experiment. **c.** 3D rendering of NM2A-eGFP fluorescent signal in a representative villus cell. **d.** Airy scan image of a live villus cell basal myosin meshwork. Surrounding neighbor cells are partially visible. Scale bar is 5 µm. **e.** Kymograph of NM2A-eGFP fluorescence, showing retrograde flow along the fiber (dashed yellow arrow). Scale bar is 10 µm. **f.** Myosin meshwork in NM2A-eGFP open-lumen organoids upon inhibiting actin retrograde flow with the cocktail of paranitro-blebbistatin, CK-666 and jasplakinolide (BCJ). Scale bar is 10 µm. **g.** Stills from a basal-plane timelapse of a NM2A-eGFP open-lumen organoid during villus cell extrusion (yellow asterisk). Images were taken every 10 s. Scale bar is 10 µm. **h.** Surface area at the basal-most plane of NM2A-eGFP extruding cells, measured at the last timepoint before fiber dissolution (‘Mesh’) and at peak fiber dissolution (‘Dissolved’). Data is from 18 extrusions, 3 independent experiments. Statistics: Wilcoxon test. **i.** PIV-vectors for the recoil of NM2A-eGFP signal after laser ablation (yellow dashed line) of the villus myosin meshwork. Scale bar is 10 µm. **j.** Extrusion of a NM2A-eGFP cell after laser-ablating the basal meshwork (yellow dashed line) in a NM2A-eGFP/mTomato mosaic open-lumen organoid. Images were taken every 10 s. **j’** XZ view of the mosaic organoid ROI after the extrusion of the NM2A-eGFP cell shown in **j** was completed. Scale bars are 10 µm. **k.** Spinning disk timelapse of a villus cell extrusion (asterisk) in open-lumen organoids expressing the Ca2+ sensor K-GECO1-FusionRed. Images were taken every 5 s. Scale bar is 10 µm. **l.** K-GECO1 Ca2+-sensor signal intensity in segmented extruding cells normalized to the mean intensity for each cell (mean +/- s.d.). x = 0 is 2.4 +/- 1.3 min (mean +/-s.e.m.) from onset of area decrease. Data is from 14 extrusions, 5 experiments. **m.** NM2A-eGFP fiber dissolution in a villus cell 2 minutes after treatment with 2 µm thapsigargin to release ER-stored calcium. Images were taken every 10 s. Scale bar is 10 µm. **n.** Immunostaining for phospho-paxillin in NM2A-eGFP open-lumen organoids in a cell in early (top) and mid (bottom) extrusion process. A cell was labelled as undergoing extrusion (yellow asterisk) if it was missing basal myosin fibers. Scale bar is 10 µm.

Fast live imaging of the basal myosin 2A meshwork showed consistent myosin 2A retrograde flow from the periphery towards the cell center in villus cells (Fig. 3e, Video 3), suggesting that this mechanism prevents myosin 2A from accumulating in the cortex of villus cells. In agreement with this interpretation, pharmacological inhibition of retrograde flow with a combination of paranitro-blebbistatin, CK-666 and jasplakinolide (BCJ cocktail^28^) abolished the fine basal meshwork (Fig. 3f, Video 4) and myosin 2A accumulated in the cell cortex. Taken together, these observations demonstrate that villus cells maintain a non-junctional, basal myosin 2A structure whose function is unknown.

We next examined whether this meshwork contributes to the villus cell extrusion process. At the onset of extrusion, the meshwork in the extruding cell rapidly dissociated to form a diffuse basal pool (Fig. 3g). This dissolution lasted around 2.5 minutes (Fig. 3g, Video 5). Simultaneously, the basal cell area expanded (Fig. 3h), indicating tension release. The presence of basal tension prior to extrusion was confirmed through laser ablation experiments on the meshwork, which showed rapid recoil away from the cut (Fig. 3i, Video 6). Within 30 minutes, 85% of cells with the meshwork damaged by laser ablation extruded from the epithelium (Fig. 3j, Video 7), indicating that tension release in the basal myosin 2A meshwork triggers extrusion. This stands in contrast to current knowledge of epithelial cell extrusion, where the first morphological change is apical constriction in the extruding cell (rev. in ^29^). Overall, these experiments identified a basal meshwork of non-junctional myosin 2A, which distinguishes villus cells from crypt cells. This meshwork sets the mechanical landscape of the villus and its dissolution initiates extrusion.

### A calcium influx precedes basal myosin meshwork disassembly

We next investigated the mechanism behind the observed rapid dissociation of the basal actomyosin meshwork. Given the known sensitivity of cytoskeletal dynamics to calcium, we hypothesized that a transient calcium influx may act as the proximal trigger of this dissociation in villus cells. To test this hypothesis, we carried out live imaging of organoids expressing a genetically encoded calcium sensor. Extrusion was preceded by a calcium pulse in the extruding cell (Fig. 3kl, Video 8), indicating that intracellular calcium might indeed be a trigger of meshwork dissolution. To directly test this, we pharmacologically elevated cytosolic calcium using thapsigargin, which led to myosin 2A meshwork dissolution within minutes (Fig. 3m, Video 9). These results, together with our finding that mechanosensitive ion channels regulate extrusion rate (Fig. 1g), suggest that villus cells extrude after an influx of calcium through mechanosensitive channels.

Because focal adhesions are necessary for basal fiber maintenance, we next examined cell-substrate adhesions in relation to basal meshwork morphology via phospho-paxillin immunostaining. In cells with a dissolved meshwork (Fig. 3n, top) as well as in those at a later stage of extrusion (Fig. 3n, bottom, Extended Data Fig. 2a), focal adhesions were indeed disassembled. In contrast, the leading edge of neighboring cells exhibited a pronounced enrichment of phospho-paxillin (Fig. 3n, bottom, arrowhead, Extended Data Fig. 2a), indicating that initial meshwork dissolution and concurrent disassembly of cell-substrate adhesions enables the protrusion of neighboring lamellipodia into the space beneath the extruding cell. Together, these experiments suggest that elevation of calcium in villus cells triggers the dissolution of the basal myosin 2A meshwork, which is the initial step in villus cell extrusion.

### The basal myosin meshwork sequesters the extrusion machinery

We next investigated how the reorganization of the myosin 2A pool following meshwork dissolution leads to extrusion. To distinguish myosin 2A dynamics in the extruding cell from that in its neighbors, we generated mosaic organoids combining membrane-tagged tdTomato (mTomato) and myosin-eGFP. Imaging the extrusion of a single myosin-eGFP cell fully surrounded by mTomato cells revealed that myosin dissolution was rapidly followed by the translocation of myosin 2A to cell-cell junctions, where it assembled into a ring (Fig. 4a). Conversely, extrusion of a mTomato cell surrounded by myosin-eGFP cells showed the formation of a second myosin ring in the neighbors. Partial mosaics (Fig. 4b, Video 10) revealed that, simultaneously with this myosin 2A reorganization, neighbor cells extended lamellipodia under the extruding cell (Fig. 4b, bottom). Altogether, these experiments reveal that in the intestine, cells simultaneously reorganize both their contractile and protrusive structures during extrusion.

**Figure 4.**
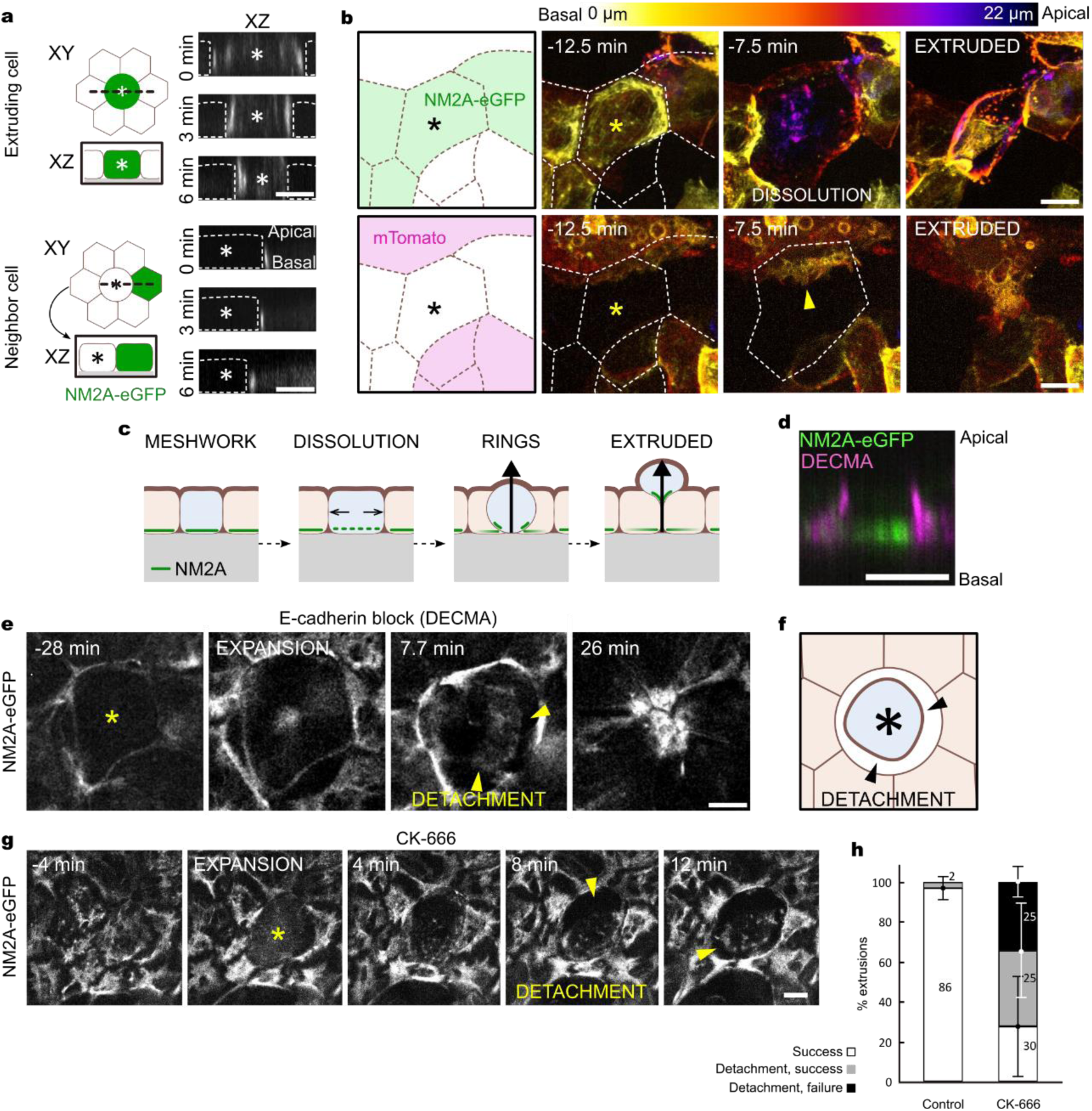
Lamellipodia ensure successful extrusion by driving the formation of the extrusion ring. **a.** Schematics and corresponding fluorescent images of mosaic open-lumen organoids showing myosin 2A ring formation in both the neighbor and extruding cell (asterisk). The NM2A-eGFP channel is shown as a reslice of the z-stack. Top: NM2A-eGFP extruding cell surrounded by mTomato cells (not shown), bottom: NM2A-eGFP neighbor cell surrounded by mTomato cells (not shown). Note the advancement of the rings towards the center of the extruding cell. Images were taken every 1.5 min. Scale bar is 10 µm. b. Left: mosaic configuration of the images shown on the right. Right: Stills from a 3D timelapse of a NM2A-eGFP/mTomato mosaic open-lumen organoid during cell extrusion of a NM2A-eGFP villus cell (yellow asterisk). Top row: NM2A-eGFP fluorescence, bottom row: mTomato fluorescence. The open arrowhead marks neighbor cell lamellipodia (bottom row). Depth of signal is color coded. Individual cells are outlined by a white dashed line. Images were taken every 2.5 min. Scale bar is 10 µm. **c.** Schematics summarizing myosin localization in relation to the villus cell extrusion sequence. **d.** E-cadherin (DECMA-1) and NM2A-eGFP apicobasal distribution in a villus cell in a fixed monolayer. Scale bar is 10 µm. **e.** Extrusion in a NM2A-eGFP open-lumen organoid treated with the E-cadherin blocking antibody (DECMA-1 clone). Images were taken every 10 s. Scale bar is 10 µm. **f.** Schematic depicting the detachment of the extruding cell from its neighbors observed mid extrusion in DECMA-treated organoids in **e**. **g.** Spinning disk timelapse of the basal-most plane of NM2A-eGFP open-lumen organoids treated with 100 µm CK-666 to inhibit actin branching and lamellipodia generation. A failed extrusion event is shown. Images were taken every 10 s. Scale bar is 10 µm. **h.** Bar plot showing the outcome of the extrusion process in control, and CK-666-treated open-lumen organoids. Data is from 3 independent experiments. Mean +/- s.d.

To assess the crosstalk between the extruding cell and its neighbors we blocked E-cadherin cell-cell connections using the DECMA function-blocking antibody (Fig. 4d, Extended Data Fig. 2b). This junction inhibition abolished the basal fibrous meshwork (Fig. 4e leftmost), indicating that intact intercellular adhesions are required to sustain meshwork tension. Instead of forming the basal meshwork, myosin 2A localized to the cell cortex. At the onset of extrusion, the extruding cell detached from its neighbor (Fig. 4ef, Video 11). Interestingly, despite the absence of E-cadherin junctions, a ring still formed in the neighboring cells and extrusion was completed. Thus, once the cell enters the extrusion process, cell-cell adhesion is necessary to form the basal myosin 2A ring in the extruding cell, but not in the neighbors. Together, these results suggest that the basal actomyosin network acts as a reservoir that sequesters the extrusion machinery until it dissolves.

### Intact lamellipodia ensure successful extrusion by driving the formation of the extrusion ring

We next asked how the dissolved myosin 2A pool translocated to cell-cell junctions to form a ring. Cryptic lamellipodial protrusions were concomitant with the formation of the myosin 2A ring in the extruding cells (Fig. 4b) and with the emergence of myosin flows towards the cell-cell junction (Video 12). These observations led us to hypothesize that cryptic lamellipodia are implicated, through E-cadherin, in the translocation of basal myosin 2A to cell-cell junctions during extrusion, thereby enabling the formation of the contractile ring in the extruding cell.

To test this hypothesis, we pharmacologically inhibited the actin-branching Arp2/3 complex with CK-666 and examined the process of ring formation. In tissues treated with CK-666 (Fig. 4g), the dissolution of the myosin 2A meshwork occurred similarly to that in control cells, indicating that lamellipodia are not required for meshwork dissolution. Instead of accumulating myosin 2A at the outer edges of the extruding cell, however, the extruding cell often detached from its neighbors (Fig. 4g, arrowheads, Video 13), similar to what we observed when we blocked E-cadherin (Fig. 4e, Video 11). In cells in which the ring-like structure formed, it was often unstable or incomplete (Extended Data Fig. 3a) and failed to fully contract around the extruding cell. Extrusion success in CK-666 treatment became dependent on cell size (Extended Data Fig. 3b): while smaller cells were still extruded efficiently, bigger ones detached and created a gap in the tissue (60% of all cells). In 30% of these cases, the cell gap continued to expand without closing (Fig. 4h, Video 13), resulting in failed extrusion and compromised tissue integrity. These results support that the interaction between cryptic lamellipodia and the overlaying extruding cells enables the formation of the ring in the extruding cell through a contact mechanism mediated by E-cadherin. This ring is dispensable for extrusion of small cells, which can be readily expelled from the tissue through forces generated by neighbors, but not for larger ones.

### Lamellipodia act as symmetry-breakers establishing apical extrusion polarity in the villus

We finally sought to link structural changes in the extruding cell and its neighbors with the generation of traction forces during extrusion. A striking feature of our spatiotemporal traction maps is that the upwards traction under the extruding cell peaks when the myosin network is dissolved and focal adhesions are disassembled. This finding led us to hypothesize that, counterintuitively, upwards tractions are generated by neighbor lamellipodia. Treatment with CK-666 abolished the peak of vertical tractions under the extruding cell (Fig. 5a), indicating that indeed lamellipodia are responsible for exerting the strong pulling-up force during extrusion. The drop in upwards traction was significant in both signed values (Fig. 5b) and magnitudes (Fig. 5c), suggesting that it results not only from an increased fraction of cells reversing the direction of vertical traction, but also from a reduction in overall traction magnitude.

**Figure 5.**
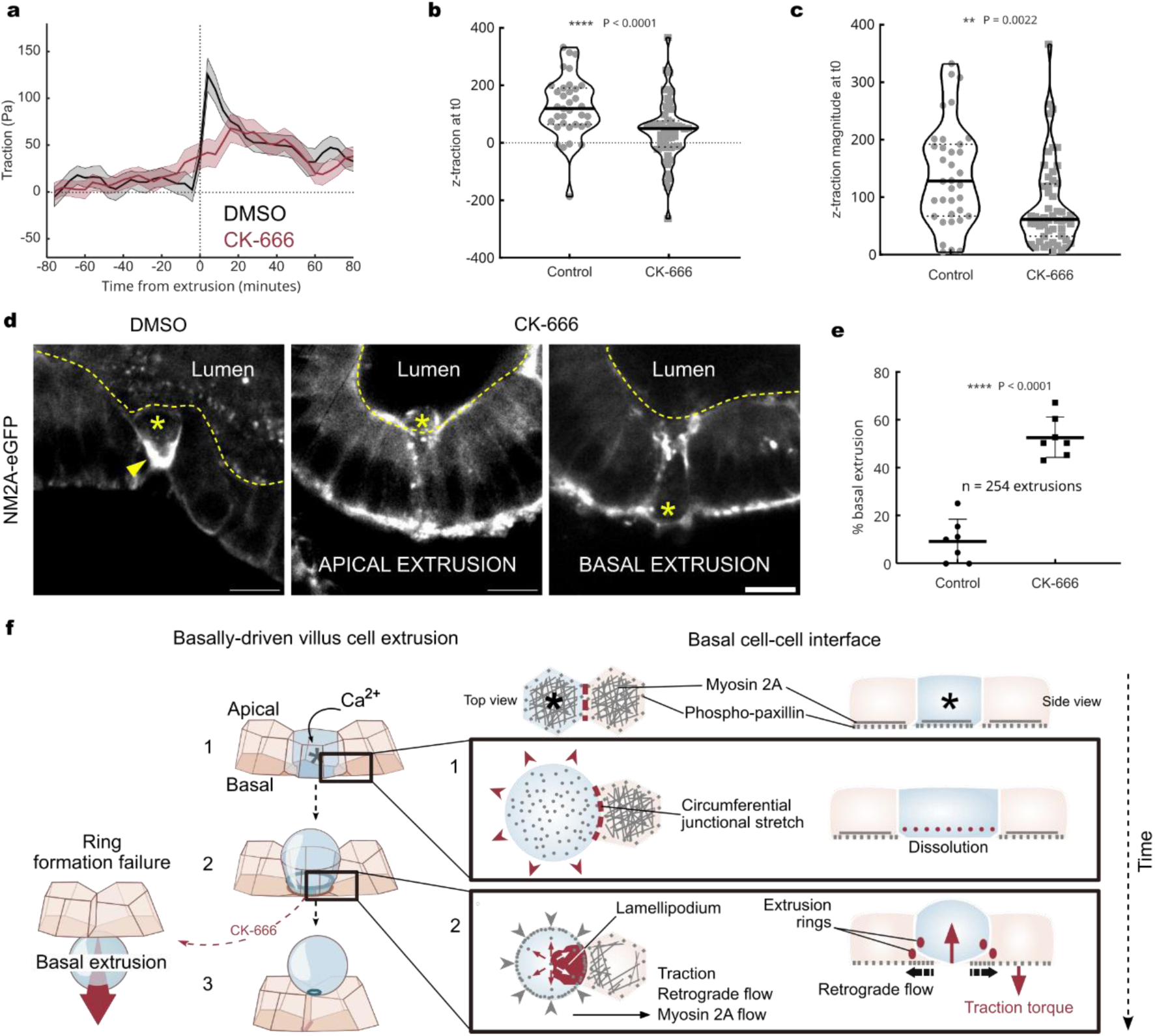
Lamellipodia act as force generators and symmetry breakers during extrusion. **a.** Average out-of-plane traction under the extruding cell (6 µm radius around cell center) as a function of time from mid extrusion, for control and CK-666-treated organoids (mean +/- s.e.m). The data is from 3 independent experiments (n=35 extrusions DMSO, n=52 extrusions CK-666). **b.** Out-of-plane traction for individual cells treated with 0.2% DMSO or 100 µm CK-666 from (a), at mid-extrusion (t0). **c.** Magnitude of out-of-plane traction for individual cells treated with 0.2% DMSO or 100 µm CK-666 from (a), at mid-extrusion (t0). **d.** Apico-basal view of extruding cells in control (leftmost) and CK-666-treated 3D NM2A-eGFP organoids. A representative example of both apical and basal (rightmost) extrusion is shown. Scale bar is 10 µm. **e.** Quantification of extrusion polarity as the fraction of basally extruding cells in 3D intestinal organoids (mean +/- s.d.). Data is from 8 individual organoids, 3 independent experiments. **f.** Schematics summarizing the proposed mechanism of basally driven villus cell extrusion. Most relevant or newly identified factors are highlighted in red.

To understand how the decrease of vertical force upon lamellipodia inhibition impacts the extrusion process, we turned to canonical 3D organoids embedded in Matrigel. In these organoids, villus regions are curved, unlike in 2D organoids, albeit in the opposite direction to that *in vivo* (Fig. 5d, yellow dashed line). When comparing control organoids with those treated with CK-666, we observed a striking difference in the intensity and localization of the basal myosin 2A during extrusion. In the control condition, myosin 2A concentrated at the basal-most plane of the extruding cell just before the cell started exiting the tissue (Fig. 5d DMSO, Video 14). This myosin 2A accumulation constricted while following the extruding cell from the basal to the apical surface (Fig. 5d DMSO, arrowhead, Video 14). By contrast, in CK-666, myosin 2A did not concentrate basally under the extruding cell, and no myosin followed the cell as it exited the tissue (Fig. 5d CK-666, Video 15). These results indicate that lamellipodia are responsible for reorganizing the villus myosin 2A meshwork into the basal ring in the extruding cell, both in 2D and 3D intestinal organoids, and independently of tissue curvature. Strikingly, in 3D organoids, lamellipodia perturbation caused cells to stop extruding exclusively into the organoid lumen. Instead, they extruded apically and basally with equal probability (Fig. 5e, Video 16). This shows that the lamellipodia-dependent basal myosin 2A ring is essential to set villus cell extrusion polarity. Together, these results show that lamellipodia are not only force generators but also symmetry breakers of intestinal cell extrusion.

## Discussion

In this study, we identified physical forces and cytoskeletal mechanisms driving cell extrusion in the mammalian intestinal epithelium. Using intestinal organoid sheets, we showed that the key event that initiates extrusion of intestinal villus cells is the sudden disruption of an actomyosin basal network and focal adhesions. Disruption is triggered by a calcium pulse and is followed by stretching of the extruding cell (Fig. 5f panel 1). This initial phase of tension relaxation is followed by the formation of new cytoskeletal structures in both the extruding cell and its neighbors: myosin 2A organizes in two contractile rings, one in the immediate neighbors and another in the extruding cell (Fig. 5f panel 2), while neighbor cells protrude cryptic lamellipodia under the extruding cell. Our data support that the formation and maintenance of these distinct structures are mechanically coupled though E-cadherin and actin retrograde flows^19,28^. Throughout the extrusion process, cells generate an upwards-pointing traction force that requires myosin but is exerted by neighbor lamellipodia. This upwards traction under extruding cells is balanced by a downwards traction in the neighbors, producing a mechanical torque that can be loosely interpreted as neighboring cells ‘shoveling out’ the extruding cell. Finally, we showed that lamellipodia are not only generators of upwards pulling tractions, but also symmetry breakers, that determine whether cells will be extruded apically or basally.

Previous studies established the role of compressive stresses in triggering extrusion, arising from tissue crowding^17^, topological defects^30^ or hypercontraction of the extruding cell itself (rev. in ^29^). In contrast, here we showed that in the intestinal epithelium, the initial phase of extrusion involves the expansion of the extruding cell, indicating that the tissue is under tension—not compression—and that part of this tension is released at extrusion onset through the dissolution of the basal actomyosin network. We also showed that curvature is not required to localize extrusion in the villus. Instead, our data suggest that the spatial distribution of myosin 2A is the main determinant of intestinal extrusion compartmentalization. Factors like crowding and curvature may enhance extrusion localization *in vivo*, particularly at the villus tip, driving the observed higher extrusion rates in this region^4,21,25,31^.

Our finding that lamellipodia are key to generating the out-of-plane extrusion force opens new fundamental questions in epithelial mechanics. Lamellipodia have not previously been considered as out-of-plane force generators in epithelia, despite evidence that they can generate subcellular torques in single cells^32^. Future studies should therefore consider their contribution to 3D epithelial mechanics, not only in extrusion but also in other functions involving out-of-plane force generation. More broadly, our study raises the question of whether the new mechanisms identified here are a distinctive feature of the intestinal epithelium or a more general strategy employed by other tissues. Besides extruding, cells in the villus must also migrate, and we previously showed that this occurs under tension gradients^24^. It is therefore appealing to consider that the intestinal epithelium has evolved a specialized extrusion mechanism that is compatible with the tensile state characteristic of actively migrating tissues^24,33^. Tissue tension thus emerges as a potential coordinator of different cellular functions, like extrusion and migration, in complex tissues composed of multiple cell types. The development of mechanically accessible organoid models, such as those presented in our study, now makes it possible to investigate the mechanobiological integration of cellular multitasking in physiologically relevant settings.

## Methods

### Organoid culture

Intestinal crypts from mice were isolated in collaborating laboratories (Matić-Vignjević laboratory and Gloerich laboratory) and used to obtain the organoid cultures from the following genetically modified mouse lines: LifeAct-eGFP^34^, mT/mG ^35^, NM2A-eGFP ^36^NM2A flox/flox-mTmG flox/flox^37^, and K-GECO1^38,39^. Organoids were passaged once a week. For passaging, the Matrigel drops (50 µl) containing organoids were mechanically disaggregated by pipetting in Phosphate-Buffered Saline (PBS) supplemented with calcium and magnesium (Sigma-Aldrich) and centrifuged at room temperature (RT) for 3.5 min at 100g. The pellet was resuspended in a 6 mg ml^-1^ Matrigel in ENR and seeded in drops of 50 µl in 24-well plates.

The ENR medium consisted of DMEM/F-12 medium (Gibco) supplemented with 2% antibiotic–antimycotic (Gibco), 2.5% GlutaMAX (Gibco), 20 ng ml^−1^ mouse EGF (Peprotech), 10 ng ml^−1^ mouse FGF (Peprotech), 100 ng ml^−1^ noggin (Peprotech), 500 ng ml^−1^ R-spondin1 (R&D Systems), 1×B-27 (Gibco) and 1×N-2 (Gibco). The Matrigel:ENR premix was kept at -20°C and placed on ice to thaw 30 min before splitting.

NM2A knockout was induced with 1 uM 4-OH tamoxifen (Sigma-Aldrich) for 24 h and induction confirmed by the presence of GFP fluorescence.

### 5 kPa Polyacrylamide (PAA) gel polymerization

35 mm glass-bottom dishes (MatTek, no. P·5G-0-20-C) were incubated for 1h at RT with Bind-silane (Sigma-Aldrich) and acetic acid (Sigma-Aldrich) dissolved in absolute ethanol (PanReac) at volume proportions of 1:1:12. Glass-bottom dishes were fully dried by aspiration. 22.5 µl of the PAA mix were added on top of the glass-bottom and covered with an 18-mm coverslip. The 5 kPa PAA mix consisted of 7.46% of Acrylamyde (BioRad), 0.044% of Bis-acrylamide (BioRad), 2% of FluoSpheres commercial solution, 0.5% of Ammonium persulphate and 0.05% of Tetramethylethylenediamine (Sigma-Aldrich). Dark red (660/680) carboxylate-modified FluoSpheres with a diameter of 0.2 µm (Thermo Fisher Scientific) were used. After exactly 1 h polymerization at room temperature (RT), gels were covered in PBS and the coverslips were removed with a scalpel blade.

### PAA gel functionalization

PAA gels were functionalized using 2 mg ml^-1^ Sulpho-SANPAH (Cultek) and irradiated with ultraviolet light (365 nm, 15 Watt) for 7.5 minutes. The gels were then washed twice with 10 mM HEPES (Gibco), followed by a quick wash with PBS, and allowed to dry at RT for 10 minutes. A 100 µl solution of 250 µg ml^−1^ rat-tail type I collagen (First Link UK) and 100 µg ml^−1^ laminin 1 (Sigma-Aldrich) dissolved in PBS was added on top of PAA gels and incubated 1 h at RT. Finally, the collagen/laminin solution was aspirated, and the gels were washed twice with 2% antibiotic–antimycotic (Gibco) PBS containing calcium and magnesium (Sigma-Aldrich).

### Cell seeding on PAA substrates

Matrigel drops containing fully grown organoids (6 days post-passage) were collected and split by pipetting in PBS containing calcium and magnesium (Sigma-Aldrich). Matrigel was removed from the mix by centrifuging at 100g for 3.5 min at RT. The 3D intestinal organoids were mechanically disaggregated again by pipetting in PBS containing calcium and magnesium (Sigma-Aldrich). For one PAA gel, we seeded the number of organoids contained in two Matrigel drops of the 24-well plate (∼200 000 cells per Matrigel drop). After disaggregation, the organoids were centrifuged at 100 g for 3.5 min at RT and the pellet was resuspended in ENR medium containing 3 µM CHIR99021 (SelleckChem), 1 mM Valproic acid (SIGMA) and 15 µM Y-27632 (Merck Millipore; "ENR-CVY" medium). The organoids were seeded in a small volume (20 µl) on top of the ECM-coated PAA gels and incubated for 1 h at 37 °C and 5% CO2. After incubation, 450 µl ENR-CVY medium was added on top of the seeded 20 µl.

For mosaic tissues, Matrigel was removed as described above. Organoids were disaggregated by pipetting with a solution containing TrypLE Express1X (Gibco) and 10 µM Y-27632 and incubated for 5 min at 37 ° C for further digestion into single cells. After digestion, TrypLE solution was inactivated by dilution and the single cell organoids were centrifuged at 200g for 5 min at RT. The pellet was resuspended in the above-mentioned medium. For one PAA gel, we seeded the number of organoids contained in four 50 µL Matrigel drops of the 24-well plate in a proportion 1:1 or 1:7 combining the two different organoid cell lines required for each specific case.

Organoids were grown in ENR medium for a minimum of 24 h prior to the start of an experiment to allow for full differentiation. All open-lumen organoid experiments were conducted 7 days post-seeding (4 days for K-GECO1 organoids), with the organoids having formed a confluent monolayer on top of the 5 kPa gel.

### 3D organoid samples

For imaging 3D organoids, cultured organoids contained in two 50 µL Matrigel drops were merged in Matrigel mixed with 0.2 µm fluorescent beads (1:50) and reseeded in 50 µL thin layers on MatTek glass-bottom dishes, following the organoid culturing protocol above. Organoids were grown in ENR medium for 5 days before imaging.

### Drug treatments

Organoid monolayers on 5 kPa gels were treated with either 0.1% DMSO, 0.2% DMSO, 50 µM zVAD-FMK (Sigma-Aldrich, cat. No. 627610-1MG), 20 µM gadolinium (III) chloride (Sigma-Aldrich, cat. No. 439770), 25 µM para-nitro blebbistatin (PN-Blebbistatin) (Motorpharma, cat. no DR-N-111), 100 µM CK-666 (Sigma-Aldrich, cat. no 182515), BCJ cocktail containing 25 µM para-nitro blebbistatin (B), 100 µM CK-666 (C) and 100 nM Jasplakinolide (J) (‘InSolution’, Calbiochem cat.no. 420127), DECMA antibody at 1:300 dilution (Sigma-Aldrich MABT26) or 2 uM thapsigargin (Thermo Fisher cat. no. T7459). Drug incubation times prior to imaging or fixation were as follows: 3 h para-nitro blebbistatin, 1 h CK-666, 1 h jasplakinolide, 72 h DECMA antibody, and no incubation (effect in 2 minutes) with thapsigargin. For treatment with the BCJ cocktail, monolayers were first incubated with para-nitro blebbistatin for 2 h, followed by the full BCJ cocktail mix for another 1 h. Incubation for the control treatments with DMSO was the same as its corresponding drug treatment. After incubation, monolayers were imaged overnight for 12 hours or immediately (thapsigargin) under the conditions listed in the Image acquisition section.

### Immunostaining

Organoid monolayers were fixed in 4% paraformaldehyde (PFA; Electron Microscopy Sciences) for 10 min at RT and washed three times with PBS. The samples were permeabilized with 1% Triton X-100 (Sigma-Aldrich) for 1 h at room temperature (RT). After three washes with washing butter (Dako, cat. No. k8007), the samples were blocked with PBS containing 0.5% Triton X-100 and 10% donkey serum (Jackson ImmunoResearch, cat. No. 017-000-121) for 1 h at RT. Primary antibodies diluted in Antibody Diluent buffer (Dako, cat. No. k8006) were added and incubated for 24 h at 4 °C. After five more washes in washing butter (5 min each), secondary antibodies and phalloidin in Antibody Diluent buffer were added for 24 h at 4 °C. Finally, the samples were washed five times with washing butter (5 min each) and three times with PBS and imaged or stored at 4 °C.

### Antibodies

Primary antibodies used and their respective dilutions were: rabbit anti cleaved caspase-3, 1:300 (Cell Signaling Technology cat. No. 9661; rabbit anti-Phospho-Paxillin, 1:200 (Cell Signaling Technology, cat. no. 69363s).

The secondary antibodies used were: goat anti-rabbit Alexa Fluor 555, 1:400 (Thermo Fisher Scientific, cat. no. A-21429); donkey anti-rabbit Alexa Fluor 647, 1:200 (Thermo Fisher Scientific, cat. no A31573). To label F-actin, phalloidin Alexa Fluor-647 (Thermo Fisher Scientific, cat. no. A22287) was used at 1:400. To label cells with active caspase 3/7 live, cells were incubated with NucView 405 (Sigma-Aldrich, cat. No. SCT104) at 3 uM for 1h prior and during imaging.

### Image acquisition

The ENR-CVY medium of the seeded organoids was replaced to normal ENR medium 24-30 h before acquisition to allow differentiation to crypts and villi. For three-dimensional traction force microscopy (TFM) experiments, images were acquired using a Nikon TiE inverted microscope with a spinning disk confocal unit (CSU-W1, Yokogawa) and a Zyla sCMOS camera (Andor). A Nikon 60X objective (Plan Apo; NA, 1.2; water immersion) was used. Specifically for TFM, the open-source Micromanager^40^ was used to carry out multidimensional acquisitions with a custom-made script. Briefly, confocal stacks of the top layer of the fluorescent beads embedded in the PAA gels were imaged with a z-step of 0.2 µm both in the deformed and relaxed states (total Z height 13 µm). Cells were imaged with a z-step of 1 µm (total Z height 20 µm). The acquisition was carried out overnight for 12 hours and images were taken every 5 minutes for each imaging position. After overnight imaging, cells were incubated inside the microscope chamber with trypsin 10X for 10 min. Samples were then washed and fluorescent beads imaged to obtain the relaxed (trypsinized) state.

For mosaic tissues, images were acquired using an Andor DragonFly 200 spinning disk confocal unit with Sona camera. A Nikon 40X objective (plan fluor; NA 0.75; dry; DIC) was used. The Fusion software was used to carry out the acquisitions. 20 µm confocal stacks with a z-step of 0.5 µm were taken in both 488 nm and 561 nm excitation channels.

For imaging a single basal tissue plane with 10 s time resolution (myosin 2A dynamics), images were acquired using the same Andor DragonFly 200 spinning disk confocal, Sona camera and a Nikon 40X air objective (plan fluor, 0.75 NA, Air, DIC) using the Fusion software. For imaging 3D organoids in Matrigel, the same imaging system and objective was used. 4D image stacks were acquired with 5 µm z-steps (total 80 µm) and 1- or 5-minute time resolution.

Time-lapse K-GECO1 images were acquired with 5 seconds time interval on a Nikon Spinning Disc confocal microscope (Yokogawa CSU-W1) using a 40X water objective (NA 1.15) in a temperature- and CO2-controlled incubator, using NIS-Elements software.

For all live experiments, a temperature box maintaining 37°C in the microscope (Life Imaging Services) and a chamber maintaining CO2 (5%) and humidity (The Brick, Life Imaging services) were used.

### Laser cuts

Laser ablation was performed using a UGA-42 Caliburn system (Rapp OptoElectronic GmbH, Germany) set at 355 nm (DPSL-355/25/TP2) with a laser power of 20% (8.4 µJ), and using a 60X Nikon objective (Plan Apo Vc 60XA/1.20 WI OFN25 DIC N2). Regions of interest were single lines with a maximum length of 10 um. These line paths were ablated through 5 repeated cuts and pulse frequency of 1 kHz. Cells for ablation were manually selected, and cuts were performed approximately 2 µm away from the outer edge of the myosin meshwork, perpendicular to the majority of the fibers. The cuts were performed in the basal plane of the monolayer.

### Image analysis

#### For cell viability

Counting extrusion events was done manually in images immunostained for cleaved caspase-3 and cell membranes labelled using the Cell Counter plugin of Fiji^41^ (ImageJ 1.54f version) Events of clear ongoing cell extrusion were manually detected in image stacks of the cell membrane fluorescence channel. An event was considered an extrusion if a cell was rounded and a part of its volume in a higher (more apical) imaging plane than its neighbors. Cells that were fully expelled from the monolayer were not considered. These cells were then manually categorized as apoptotic if (cleaved) caspase-3 antibody signal was higher than background signal.

#### For extrusion rate

Counting of extrusion events was done manually in timelapse images (4- or 5-min time resolution) with cell membranes or non-muscle myosin 2A labelled, using the Cell Counter plugin of Fiji. Then, all cells in the field of view were manually counted using the same tool, in the first and in the last timepoint. Crypt regions were segmented manually using the Fiji Polygon Selections tool based on the recognizable circular crypt morphology, including the basally constricted transit-amplifying region. Based on these segmentation masks, cells were categorized into crypt or villus cells. The number of cells in the monolayer was calculated as the average number of cells in the first and in the last timepoint images. Extrusion rate (% of cells/h) was then obtained by dividing the total number of counted extrusion events, by the number of cells in the monolayer and the number of hours imaged (minimum 6 h).

#### For extrusion rate spatial distribution

Using a custom script (MATLAB), crypt areas were linearly interpolated between the first and the last timepoint to obtain an estimate of crypt edge. Similarly, cell positions were linearly interpolated between the first and last timepoint, along a linear track obtained from the TrackMate Fiji plugin. Both crypt edges and cell positions were marked as described in the previous section. In each timepoint image, the field of view was divided into 20 µm-wide radial bins, defined by distance from crypt edge. In fields of view that contained more than one crypt, distance bins were defined for each crypt edge. The 20 µm bin width was chosen in order to include at least one average cell diameter (14 µm). Then, for each extruding cell, its closest (‘mother’) crypt was identified, and this cell was assigned a distance bin based on the Euclidean distance to its mother crypt. After this, the extrusion rate was calculated for each bin as described in the previous section. Finally, both quantities were averaged over all time points.

#### For myosin intensity in crypt and villus

Using Fiji, crypts were segmented in basal planes of NM2A-GFP organoid image stacks (0.5 µm z-step) as described above (‘*For extrusion rate.*’). Villus or crypt images were generated by removing crypt, or non-crypt areas, respectively, from original image stacks. To obtain the total amount of myosin (GFP) signal in the villus or the crypt, total signal intensity was first projected into a single stack using the Fiji sum projection and then all pixel values were summed up using the raw integrated density measurement. This total signal was then divided by the total number of crypt or villus cells to obtain the average signal intensity per cell, per tissue compartment.

#### For three-dimensional traction force microscopy

Data consisted of a collection of Z-stacks of the cell-substrate interphase at different timepoints (taken every 5 minutes for 12 hours as described above). To measure traction forces under a cell before extrusion, cell back-tracking of the extruded cells was done using Fiji. The Fiji analysis included manually identifying the extrusion events, segmenting the extruding cell by drawing a polygon at the basal-most plane. The centroid, along with other morphological parameters of the segmented cell was measured. The extruding cell was backtracked and segmented in this way at several morphological hallmark timepoints. Cell shape in between these timepoints was interpolated using Fiji’s ‘Interpolate ROIs’ tool. Each extruding cell was backtracked for 30 time points (150 min) before the extrusion event. Extrusion timepoint (t0) was defined as the last timepoint when the extruding cell was visible at the basal-most tissue plane.

The 3D tractions were computed as previously described^42^. Briefly, the displacement field was obtained from fluorescent bead images. Tractions were calculated from displacements. Traction vectors were decomposed in x, y and z and the component of the 3D vector pointing towards the centroid of the tracked cell was used for further analyses of radial tractions and the vector z component for analyses of vertical tractions.

#### For optical flow

To compute the recoil velocities, PIV was performed using a window size of 32 × 32 pixels and an overlap of 0.75 using in-house code.

### Data analysis and statistics

Data was analyzed using MATLAB 2020b or ImageJ 1.54f macros. Graphs were plotted using MATLAB, PRISM 8.0.2 or Excel and edited for style using Inkscape. Data was tested for statistical significance using PRISM 8.0.2 with the t-test if normality was met or could not be tested (paired if individual measurements were the biggest source of variability, all except Fig. 5; or unpaired in Fig. 5), or the Mann-Whitney test or Wilcoxon test when normality test was not passed, depending is the data was paired or not, respectively. Differences between conditions were considered statistically significant when p < 0.05.

## Acknowledgements

We thank Danijela Matić-Vignjević for providing organoids and Reda Bouras for technical help with preparing organoids. We thank all the members of the Roca-Cusachs and Trepat laboratories, as well as Marino Arroyo, for their discussions and support. Funding: M.M. acknowledges funding from the Spanish Ministry for Science and Innovation (Juan de la Cierva FJC2018-037440-I) and EMBO (ALTF-1169). P.R.-C. acknowledges funding from the Spanish Ministry of Science and Innovation (PID2022-142672NB-I00), the European Research Council (AdG 101097753), the Generalitat de Catalunya (2017-SGR-1602), and the prize ‘ICREA Academia’ for excellence in research. X.T. acknowledges funding from the Generalitat de Catalunya (AGAUR SGR-2017-01602), the CERCA Programme, the Spanish Ministry for Science and Innovation MICCINN/FEDER (PID2021-128635NB-I00 MCIN/AEI/ 10.13039/501100011033 and “ERDF-EU A way of making Europe”), European Research Council (Adv-883739), Fundació la Marató de TV3 (project 201903-30-31-32), European Commission (H2020-FETPROACT-01-2016-731957), La Caixa Foundation (LCF/PR/HR24/00326) and the Human Frontiers Science Program (HFSPRGP022/2024); IBEC is recipient of a Severo Ochoa Award of Excellence from the MINECO.

## Author contributions

M.M and X.T. conceived the project. M.M. designed and performed the experiments, wrote analysis tools and analyzed the data. M.W. and E.L.S. helped with experiments and with cell tracking. C.P.G wrote tractions-related extrusion analysis tools. C.P.G and G.C. contributed microscopy images. R.H. performed experiments with the K-GECO1 organoid line. P.R.C. and M.G. provided material and discussion. M.M. and X.T. wrote the manuscript. All authors revised the completed manuscript.

## Code availability

Analysis procedures and code are available from the corresponding authors upon reasonable request.

## Data availability

The data that support the findings of this study are available from the corresponding authors upon reasonable request.

## Additional information

Extended Data is available for this paper.

Correspondence should be addressed to M.M. or X.T. and requests for materials to X.T.

