## Extended Data for "Mechanical Coordination of Intestinal Cell Extrusion by Supracellular 3D Force Patterns"


This file contains:

Extended Data Figures

Supplementary Video Legends

### Extended Data Figures

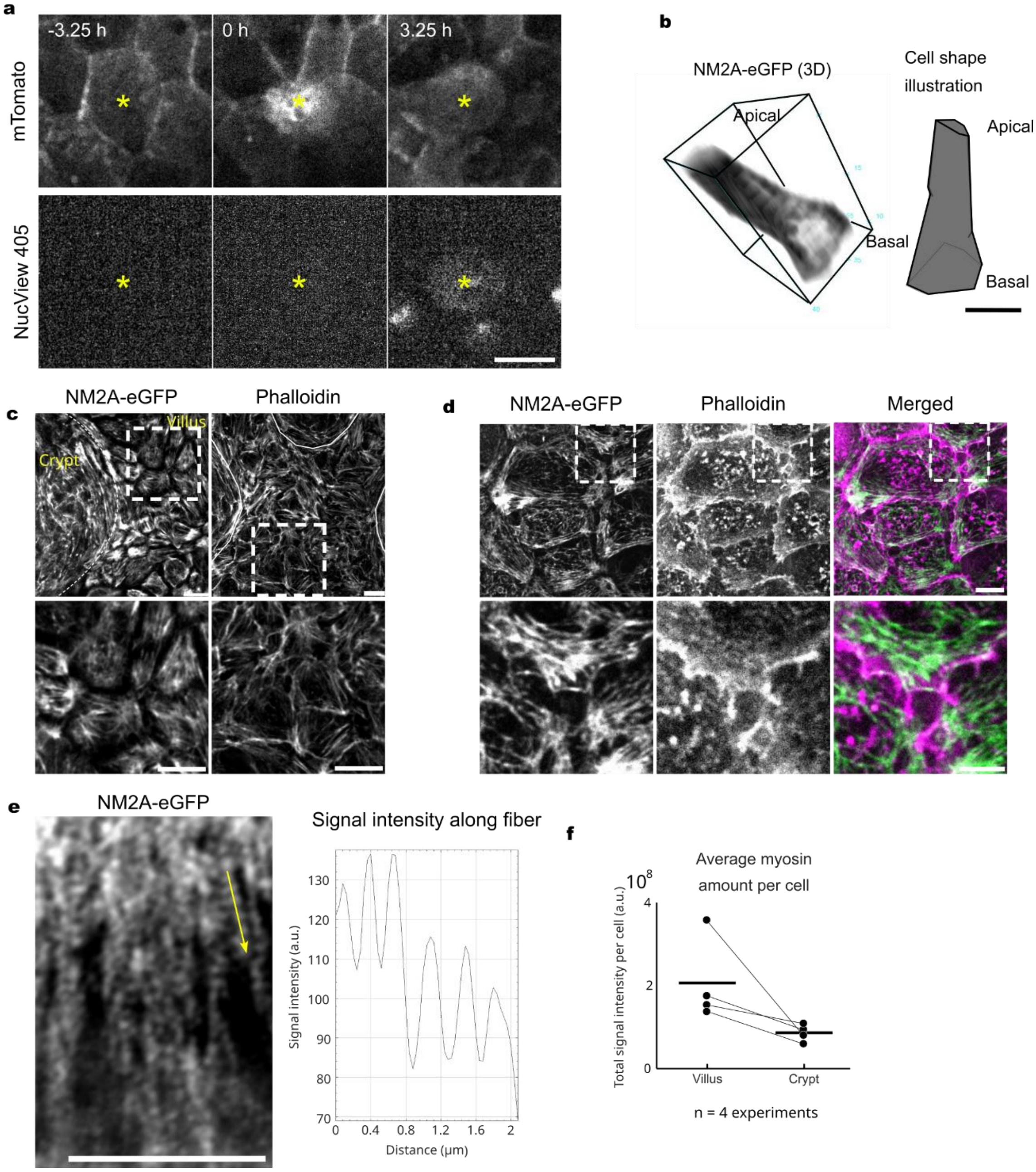


**Extended Data Fig. 1. Villus cells do not require apoptosis to extrude and are distinguished from crypt cells by a basal myosin 2A meshwork. a.** Spinning disk image of a cell extruding from an mTomato open-lumen organoid treated with 3 μM NucView 405 to label active caspase 3/7. Scale bar: 10 μm. **b.** 3D rendering (Fiji 3D Viewer) of NM2A-eGFP signal in a typical crypt cell (right). Illustration of the segmented cell shape for the same cell. Scale bar: 5 μm. **c.** PFA-fixed NM2A-eGFP open-lumen organoid tissue stained for Phalloidin to label F-actin. Bottom row: enlarged image of the dashed regions in the top row. Scale bars: 10 μm. **d.** Basal plane of the villus region of PFA-fixed NM2A-eGFP open-lumen organoid tissue stained for Phalloidin to label F-actin. Bottom row: enlarged image of the dashed regions in the top row to focus on the junctional exclusion of myosin 2A. Scale bars: 10 μm (top), 5 μm (bottom). **e.** Right: Airy scan image of a peripheral part of the basal myosin meshwork in a NM2A-eGFP cell. Left: NM2A-eGFP fluorescence signal intensity along line marked in the left image (yellow arrow). Scale bar: 5 µm. **f.** Average amount of NM2A-eGFP fluorescence signal integrated over the z-stack of the villus or crypt compartment. Data from 4 independent experiments.

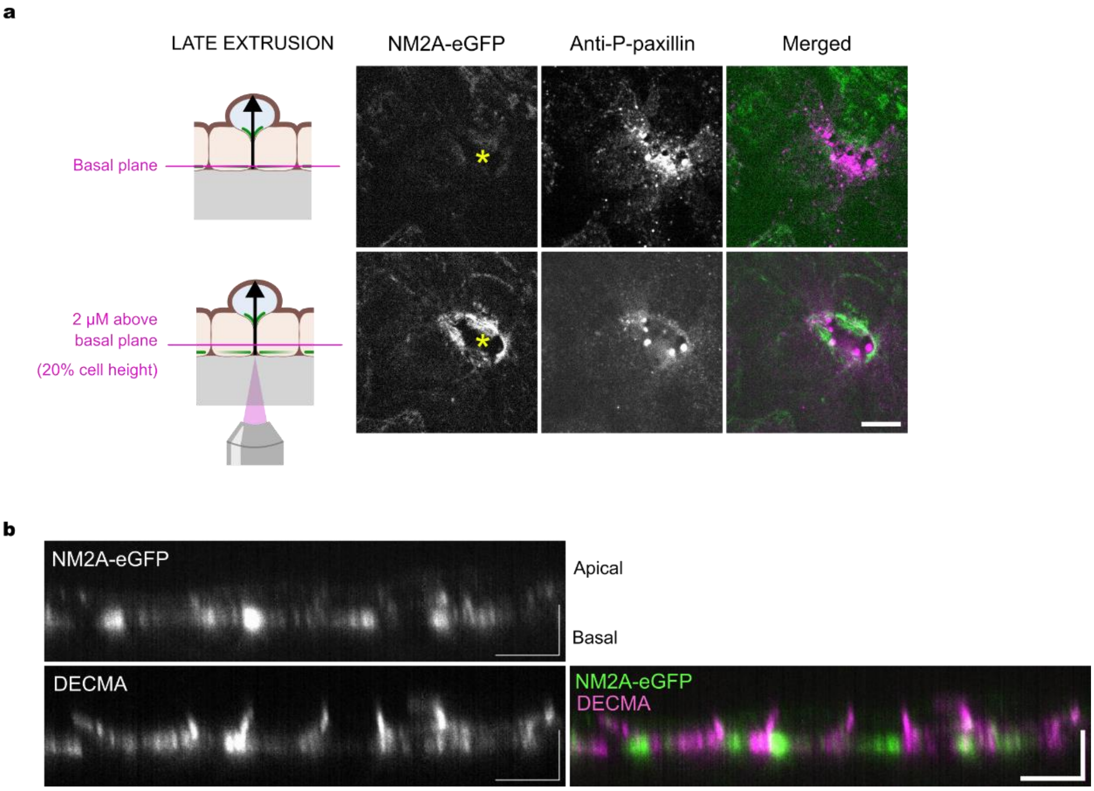


**Extended Data Fig. 2. Focal adhesions during extrusion and E-cadherin adhesions in open lumen organoids. a.** Immunostaining for phospho-paxillin in NM2A-eGFP open-lumen organoids in a cell in the late stage of the extrusion process at the most basal plane (top) and 2 µm above it (bottom). A cell was labelled as undergoing extrusion (yellow asterisk) if it was missing basal myosin fibers. Scale bar is 10 µm. **b.** Apico-basal view of the immunostaining for E-cadherin (DECMA-1 clone) in NM2A-eGFP open-lumen organoids. Note the basolateral distribution of E-cadherin (magenta). Scale bars: 10 µm.

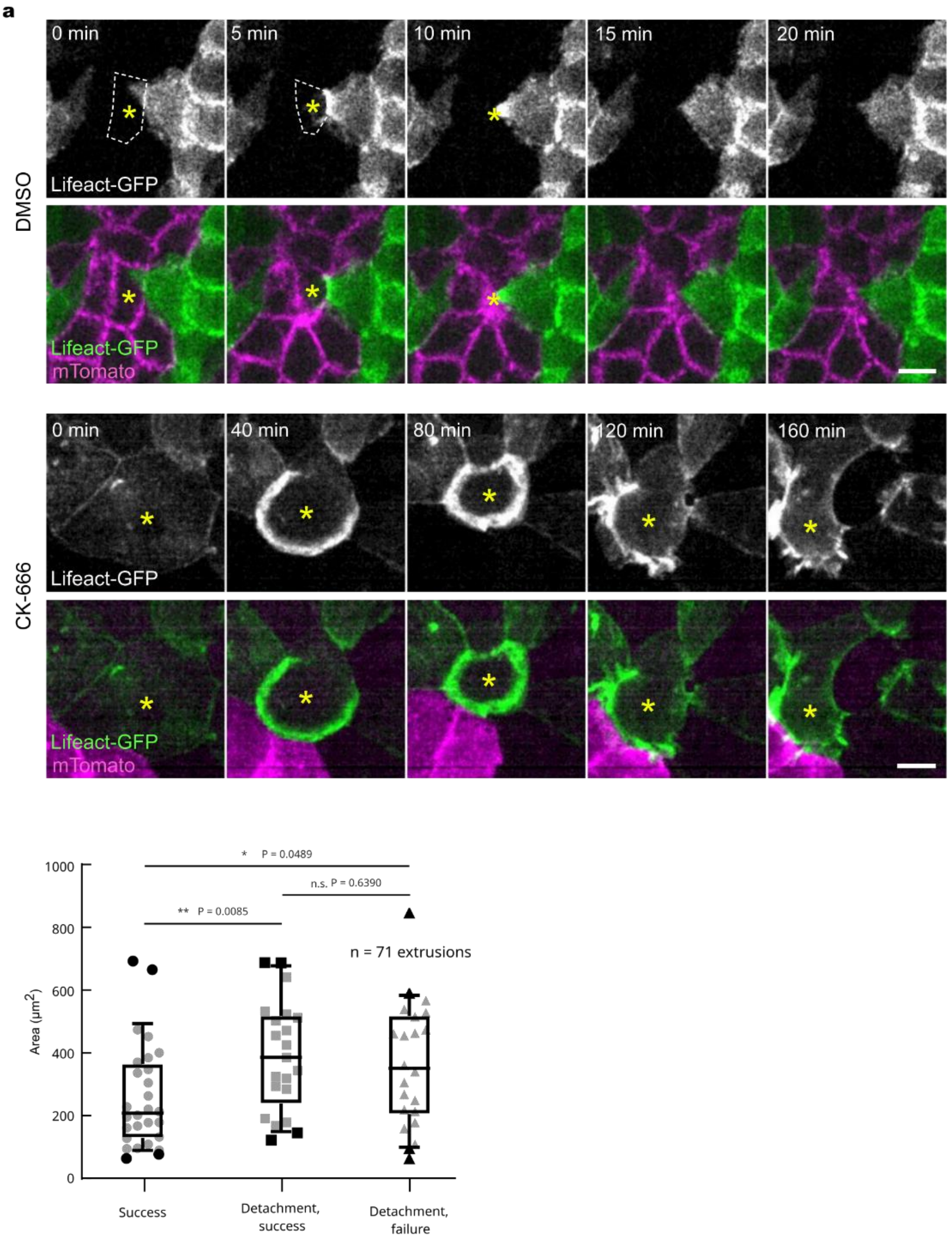


**Extended Data Fig. 3. Perturbation of lamellipodia causes ring formation failure and cell size-dependent extrusion failure. a.** Top: Generation of the actin ring in control mosaic open-lumen organoids. Bottom: Generation and failure of the actin ring in the extruding cell in CK-666 treated mosaic open-lumen organoids. Note the longer timescale of the process compared to DMSO above. Scale bars: 10 µm. **b.** Area of extruding cells in CK-666-treated open-lumen organoids in relation to the outcome of the extrusion process (success, detached cells but successful extrusion, or failure). Areas were segmented at the basal cell plane in the last timepoint (10 s) before extrusion entry point, i.e., meshwork dissolution or cell area expansion. Statistics: Mann-Whitney. Data is from 3 independent experiments. Cell number per category: 28, 21, 22.

#### Supplementary Video Legends

**Supplementary Video 1. Villus cell extrusion in open-lumen organoids.**
Spinning-disk timelapse of an extrusion of a villus cell (yellow asterisk). Cell membranes are labelled with membrane-tagged tdTomato (mTomato). Images were taken every 4 min.

**Supplementary Video 2. TFM during villus cell extrusion in open-lumen organoids.**
XZ-view of a spinning disk timelapse of an intestinal extrusion event, with traction vectors measured with traction force microscopy (yellow arrows). Images were taken every 4 min. Traction scale bar is 100 Pa.

**Supplementary Video 3. Retrograde flow.**
Spinning disk timelapse of NM2A-eGFP fluorescence, showing retrograde flow from the cell periphery towards the cell center. Images were taken every 10 s.

**Supplementary Video 4. Retrograde flow inhibition.**
Spinning disk timelapse of the myosin meshwork dynamics in NM2A-eGFP open-lumen organoids upon inhibiting actin retrograde flow with the cocktail of paranitro-blebbistatin, CK-666 and jasplakinolide (BCJ). Images were taken every 10 s.

**Supplementary Video 5. Myosin meshwork reorganization during extrusion.**
Basal-plane timelapse of a NM2A-eGFP open-lumen organoid during villus cell extrusion (yellow asterisk). Images were taken every 10 s.

**Supplementary Video 6. Laser ablation of the myosin meshwork fibers.**
Recoil of NM2A-eGFP signal after laser ablation of the basal myosin meshwork in a villus cell. Images were taken every 150 ms.

**Supplementary Video 7. Cell extrusion after laser ablation of the myosin meshwork.**
Spinning disk timelapse of extrusion of a NM2A-eGFP cell after laser-ablating the basal meshwork in a NM2A-eGFP/mTomato mosaic open-lumen organoid. Images were taken every 10 s.

**Supplementary Video 8. Calcium during villus cell extrusion.**
Spinning disk timelapse of a villus cell extrusion (yellow asterisk) in open-lumen organoids expressing the $Ca^{2+}$ sensor KGECO1-FusionRed. Images were taken every 5 s.

**Supplementary Video 9. Thapsigargin treatment.**
Spinning disk timelapse of NM2A-eGFP fiber dissolution in a villus cell 2 minutes after treatment with 2 µm thapsigargin to release ER-stored calcium. Images were taken every 10 s.

**Supplementary Video 10. Lamellipodial protrusions during extrusion in a mosaic open-lumen organoid.**

Spinning disk timelapse of mTomato neighbor cells during extrusion of a NM2A-eGFP cell (only outlines shown) in a NM2A-eGFP/mTomato mosaic open-lumen organoid. Images were taken every 2.5 min.

**Supplementary Video 11. Extrusion under E-cadherin block.**
Spinning disk timelapse of cell extrusion in a NM2A-eGFP open-lumen organoid treated with the E-cadherin blocking antibody (DECMA-1 clone). Note that a myosin ring forms only in the immediate neighbors and fails to form in the extruding cell. Images were taken every 10 s.

**Supplementary Video 12. Myosin 2A flow to junction during extrusion.**
Spinning disk timelapse showing the relocalization of the NM2A-eGFP protein from non-junctional to junctional after damaging the basal villus cell meshwork with laser ablation (yellow line) in cell A. Images were taken every 5 s.

**Supplementary Video 13. Failed extrusion in CK-666.**
Spinning disk timelapse of a failed extrusion in a NM2A-eGFP open-lumen organoid treated with 100 µm CK-666. Images were taken every 10 s.

**Supplementary Video 14. Extrusion in 3D organoids.**
Spinning disk timelapse of the apico-basal view of an extruding cell in a control 3D NM2A-eGFP organoid. Images were taken every 1 min.

**Supplementary Video 15. Extrusion in 3D organoids treated with CK-666.**
Spinning disk timelapse of the apico-basal view of an extruding cell in a 3D NM2A-eGFP organoid treated with CK-666. Images were taken every 1 min.

**Supplementary Video 16. Basal extrusion in 3D organoids treated with CK-666.**
Spinning disk timelapse of the apico-basal view of a basally extruding cell in a 3D NM2A-eGFP organoid treated with CK-666. Images were taken every 1 min.
